# Understanding the Mechanistic Contribution of Herbal Extracts in Compound Kushen Injection with Transcriptome Analysis

**DOI:** 10.1101/592964

**Authors:** Hanyuan Shen, Zhipeng Qu, Yuka Harata-Lee, Thazin Nwe Aung, Jian Cui, Wei Wang, R. Daniel Kortschak, David L. Adelson

## Abstract

Herbal compatibility is the knowledge of which herbs to combine in traditional Chinese medicine (TCM) formulations. The lack of understanding of herbal compatibility is one of the key problems for the application and popularization of TCM in western society. Because of the chemical complexity of herbal medicines, it is simpler to begin to conduct compatibility research based on herbs rather than component plant secondary metabolites. We have used transcriptome analysis to explore the effects and interactions of two plant extracts (Kushen and Baituling) combined in Compound Kushen Injection (CKI). Based on shared chemical compounds and *in vitro* cytotoxicity comparisons, we found that both the major compounds in CKI, and the cytotoxicity effects of CKI were mainly derived from the extract of Kushen (*Sophorae flavescentis*). We generated and analyzed transcriptome data from MDA-MB-231 cells treated with single-herb extracts or CKI and results showed that Kushen contributed to the perturbation of the majority of cytotoxicity/cancer related pathways in CKI such as cell cycle and DNA replication. We also found that Baituling (*Heterosmilax yunnanensis Gagnep*) could not only enhance the cytotoxic effects of Kushen in CKI, but also activate immune-related pathways. Our analyses predicted that IL-1**β** gene expression was upregulated by Baituling in CKI and we confirmed that IL-1**β** protein expression was increased using an ELISA assay. Altogether, these findings help to explain the rationale for combining Kushen and Baituling in CKI, and transcriptome analysis using single herb extracts is an effective method for understanding herbal compatibility in TCM.

## Introduction

At present many complex and chronic diseases rely on therapies that combine modern pharmaceuticals. A similar multiple-herb strategy known as “Fufang” is an essential component in traditional Chinese medicine (TCM) theory and is used to achieve better therapeutic results, and reduce side effects and herbal toxicity^1–2^. As result of thousands of years’ of accumulated clinical practice, TCM has more than 100,000 formulae and abundant experience that has contributed to an understanding of which herbs should be combined in particular circumstances, herein referred to as herbal compatibility^3,4^. However, because herbal medicines are made up of complex mixtures of plant secondary metabolites, the mechanism of most TCM formulas has not been explored. This limitation of TCM has become one of the key problems for its modernization, and hinders the application and popularization of herbal medicines^5^.

Recent rapid developments in analytical chemistry and molecular biology have provided methods for researchers to tackle the complex mechanisms of herbal compatibility on a number of different levels. Usually, these methods focus on one or a few components within a complex mixture, in attempts to reveal how preparation/extraction for combined use can change their concentrations in products or pharmacokinetic processes *in vivo*^*6–9*^. However, as a complex mixture may contain thousands of compounds, it is unclear how changing one or several components in a TCM formula can explain and account for the principles and observations of herbal compatibility. Furthermore, pharmacological models that measure phenotypes associated with efficacy or proxies for efficacy are limited in their ability to explain potential therapeutic effects and mechanisms. New high-throughput technologies for measuring molecular phenotypes such as gene expression, and bioinformatic methods can provide systematic ways for refining and clarifying complex biological processes that result from hundreds or thousands of molecular interactions. By applying these methods to the study of TCM, it is possible to transform the research paradigm from “main active compound that influences one target” to “multiple components that influence many network targets”^10,11^. Although it is now common to apply RNA-sequencing and systematic methods to study the effects of whole TCM formulae, no literature has applied these methods to study herbal compatibility. In this report, we apply transcriptome analysis to identify how the combination of Kushen (*Sophorae flavescentis*) and Baituling (*Heterosmilax yunnanensis Gagnep*) extracts can account for the broader and increased effects observed in Compound Kushen Injection (CKI).

Our model system for dissecting herbal compatibility, CKI, is derived from an ancient Chinese formula and was approved by the State Food and Drug Administration (SFDA) of China in 1995. It is widely used as an adjuvant medicine in the treatment of carcinomas for pain relief, activation of innate immune response and reduced side effects in cancer therapy^12,13^. Our previous results have shown that CKI suppresses the growth of cancer cells by inhibiting cell cycle, energy metabolism, and DNA repair pathways^11,14^. Kushen is considered to be the principal herb and major contributor to the molecular effects of CKI. Many published studies have reported on the alkaloids and flavonoids contained in CKI, most of which are extracted from Kushen. These compounds have been reported to have a variety of bioactivities, including antitumor, antioxidant and anti-inflammatory activities^15,16^. However, there is no literature that mentions the role of Baituling in CKI. Therefore, current studies are not sufficient to provide a rational framework for CKI prescription or explain its molecular mechanisms.

In this report, we break down the formula into its individual components in order to study the herbal compatibility of CKI. By comparing transcriptome changes in MDA-MB-231 cells between CKI and single herbal extract treatments, we found that Kushen extract alone, perturbed most of the pathways through which CKI exerts its effects on cancer cells. However, integrating Kushen with Baituling can enhance the effects of Kushen alone on cancer-related pathways and in addition can activate innate immune functions. These results support the TCM rationale behind the CKI formula and confirm RNA-sequencing as a useful tool for the identification of candidate mechanisms in TCM research.

## Results

### HPLC comparison of the composition of CKI and single extract injections

In order to obtain information about plant specific components in CKI, we used high-performance liquid chromatography (HPLC) to compare the compounds in CKI and two single herb extracts/injections (Fig.1). From the chromatographic profile, it can be seen that Kushen injection contributes most of the major chemical components in CKI. In contrast, few compounds were detected in Baituling injection, which only contributes one major compound to CKI. Based on comparison with 9 reference standard compounds, 8 main compounds derived from Kushen (adenine, N-methylcytisine, sophorodine, matrine, sophocarpine, oxysophocarpine, oxymatrine and trifolirhizin) are shown to contribute to CKI. Macrozamin, which is used as a control marker for Baituling during manufacturing, only appeared in CKI and Baituling injection. These results indicate that CKI contains major compounds from both Kushen and Baituling, and Kushen contributes most of the major chemical components in CKI.

**Figure 1:**
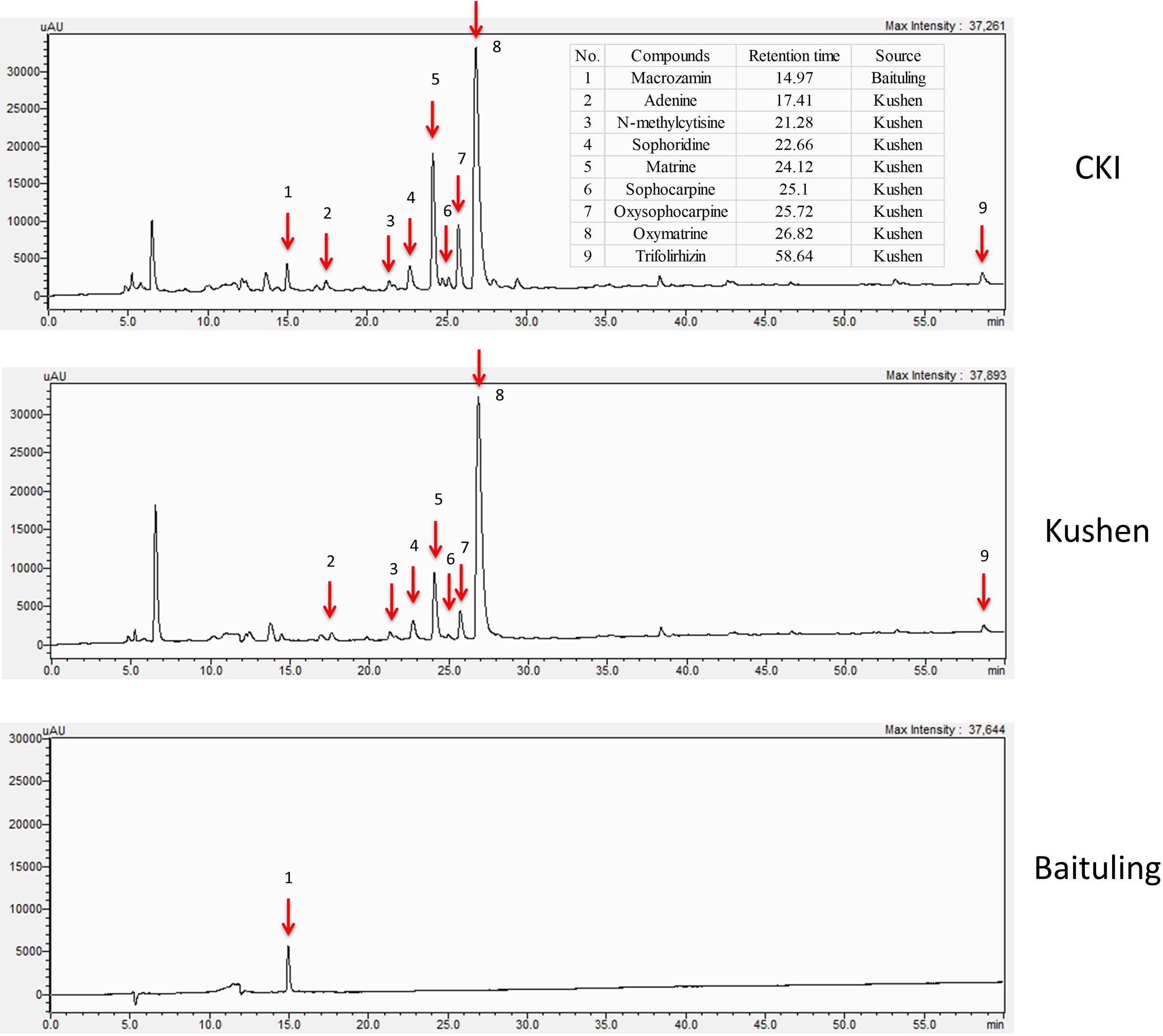
HPLC profiles of CKI, Kushen and Baituling injections. Nine component compounds characterized using standard compounds are marked with red arrows.

### Comparison of the anticancer effects between CKI and single injections

Our previous results showed CKI suppressed the proliferation of and induced apoptosis in MCF-7 cells^11^. To determine whether single injections had similar phenotypic effects as CKI, we conducted XTT assays to measure cell viability using three different cell lines; MDA-MB-231, A431 and HepG2. Results showed that Kushen had stronger cytotoxic effects than Baituling in all three cell lines. However, neither of the single injections had apoptotic effects comparable to CKI (Fig.2A) based on rates of apoptosis determined by flow cytometry with propidium iodide (PI) staining. Consistent with XTT cell viability results, more apoptotic cells were found in CKI treatments than single injections and BTL had the smallest effect on apoptosis of the three injections (Fig.2B).

**Figure 2:**
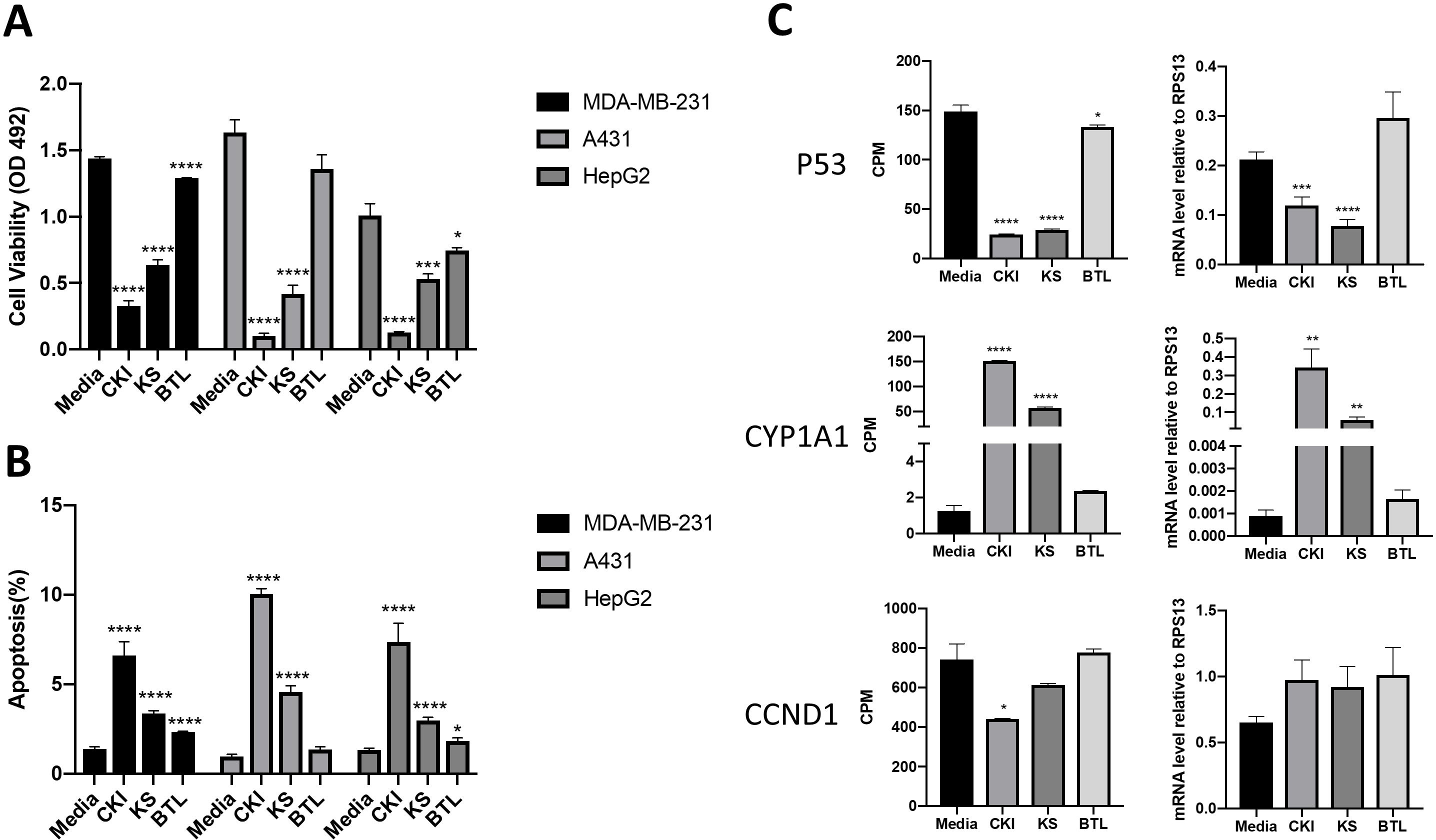
Comparison of the effect of CKI and single herb injections on cancer cell lines. A. And B. Viability and the percentage of apoptotic cells treated with different injections for 48 hours. C.q-PCR validation of RNA sequencing results. Results are represented as means ±SEM (n=9). Statistical analyses were performed using the t-test compared to untreated “Media” (*p<0.05, **p<0.01, ***p<0.001, ****p<0.0001)

### Comparison of MDA-MB-231 transcriptomes from CKI and single injection treatments

In order to elucidate the molecular mechanisms of herbal compatibility in CKI, we carried out transcriptome profiling from CKI and single injection treated MDA-MB-231 cells. Triplicate samples for each treatment clustered well in multidimensional scaling plots and different treatments were clearly separated (Supplementary Fig 1). Although CKI and Kushen injection have similar chemical profiles, the inclusion of Baituling in CKI is sufficient to change the transcriptome of MDA-MB-231 cells compared to Kushen single injection treatment. We used edgeR^17^ to identify differentially expressed (DE) genes for each injection treatment compared to untreated. In addition, we also identified DE genes for CKI treatment compared to Kushen treatment to identify the effects of Baituling in CKI (Supplementary Table 3).

To validate the results of transcriptome analysis, we performed quantitative PCR for several genes known to be important read-outs for the effects of CKI; TP53, CYD1A1 and CCND1. Their expression levels confirmed the interaction of Baituling with Kushen observed in the overall RNA sequencing results (Fig.2C).

### Similar effects of CKI and Kushen single injection on MDA-MB-231 cells

Because Kushen is considered to be the primary active herb in CKI, we first examined the effects of Kushen and CKI to see if they had similar effects on genes. By comparing the DE gene set between Kushen and CKI, the shared DE gene group accounted for 63% and 81% of DE genes from CKI and Kushen respectively (Fig.3A), indicating that Kushen contributed to the majority of effects from CKI.

**Figure 3:**
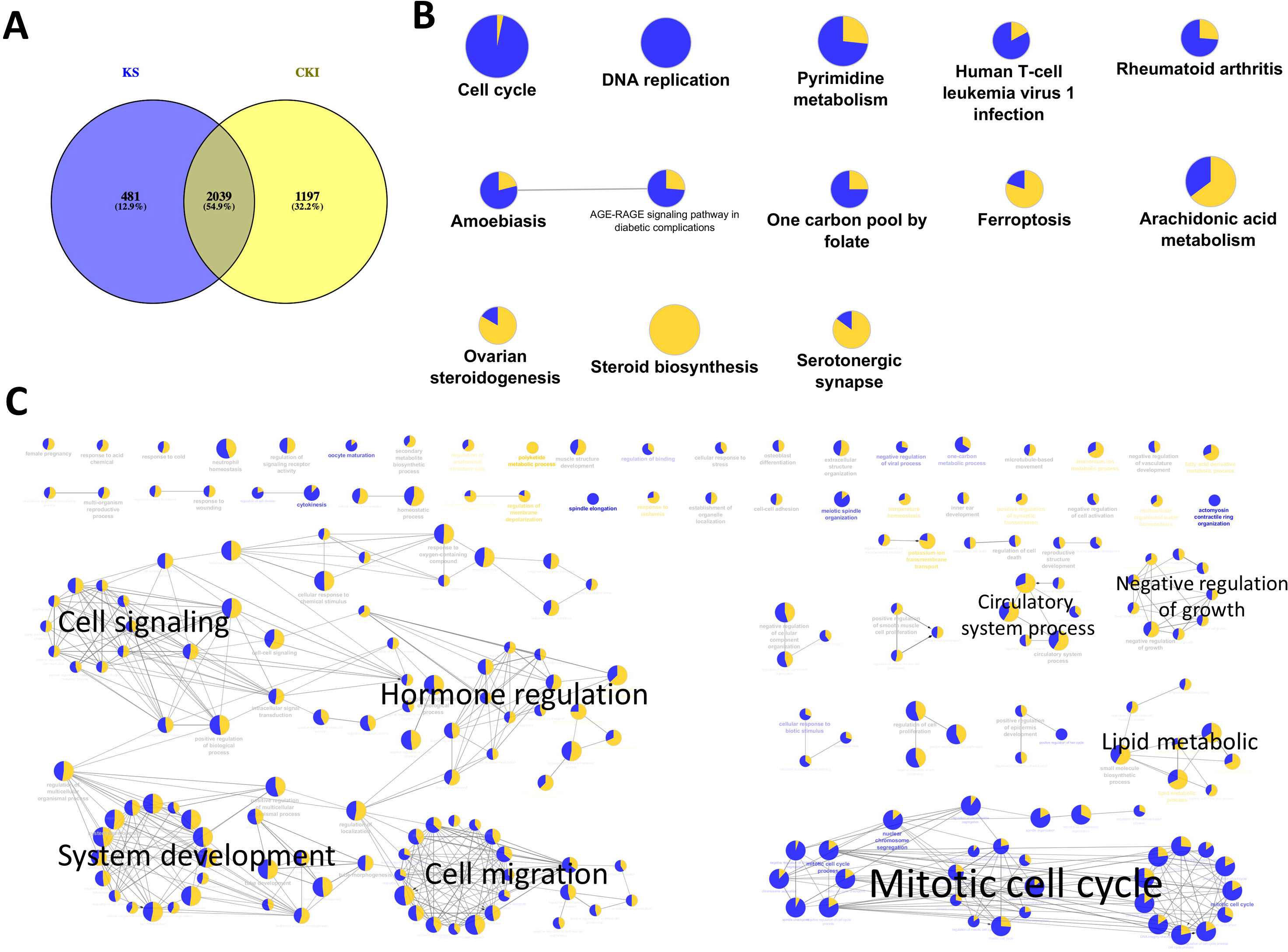
Significantly differentially expressed genes shared by Kushen and CKI treated cells and their functional enrichment analysis. A. Venn diagram showing the number of differentially regulated genes in MDA-MB-231 cells treated with Kushen (KS; blue) or CKI (yellow) compared to untreated cells. B. and C. Over-represented KEGG pathways and GO terms (Biological Process at 3rd level) for genes similarly regulated by Kushen and CKI. Node size is proportional to the statistical significance of over-representation and colors represent the proportion of up or down-regulated genes (yellow=up-regulated and blue=down-regulated). Similar GO terms are clustered (with representative terms shown in bold) and connected with edges.

In order to better understand the functions of shared DE genes between Kushen and CKI, we performed over-representation analysis using Gene Ontology (GO) and Kyoto Encyclopedia of Genes and Genomes (KEGG) pathways for these 2039 genes(Fig.3B&C, Supplementary Table 5). The results showed that cell cycle- and DNA replication-related pathways and terms were largely down-regulated by both Kushen and CKI. Based on our previous publications, perturbed regulation of genes in these pathways and annotated by these terms was associated with the observed cell viability and apoptotic effects CKI on cancer cells^11,14^. Other terms showing perturbation of metabolic processes and cell migration identified in this study, such as ‘pyrimidine metabolism’, ‘steroid biosynthesis’ and ‘positive regulation of locomotion’, also showed up in our previous research on the effects of CKI. Altogether, these results indicated that Kushen was very important to the major molecular consequences of CKI treatment, and perturbed most of the functions perturbed by CKI, including reduced viability and apoptosis in cancer cells.

### Different effects of CKI and single injections on MDA-MB-231 cells

Although the above results for enrichment analysis showed that Kushen and CKI mainly regulate the same pathways, they did not specifically show the magnitude and direction of these perturbations. We, therefore, performed Signalling Pathway Impact Analysis (SPIA) to compare the significantly perturbed functional pathways across the different treatments. Ninety two pathways were found to be significantly perturbed by CKI while only 30 of them were shown to be activated. For Kushen and Baituling, the ratios of activated/inhibited were 29/100 and 24/58 respectively (Supplementary Table 4). Clearly, Baituling perturbed fewer pathways, but the ratio of activated/inhibited was higher than for Kushen or CKI. Interestingly, most pathways were perturbed in the same way (inhibited or activated) by CKI and Kushen as shown in Fig.4A, with very few pathways showing different types of perturbation.

**Figure 4:**
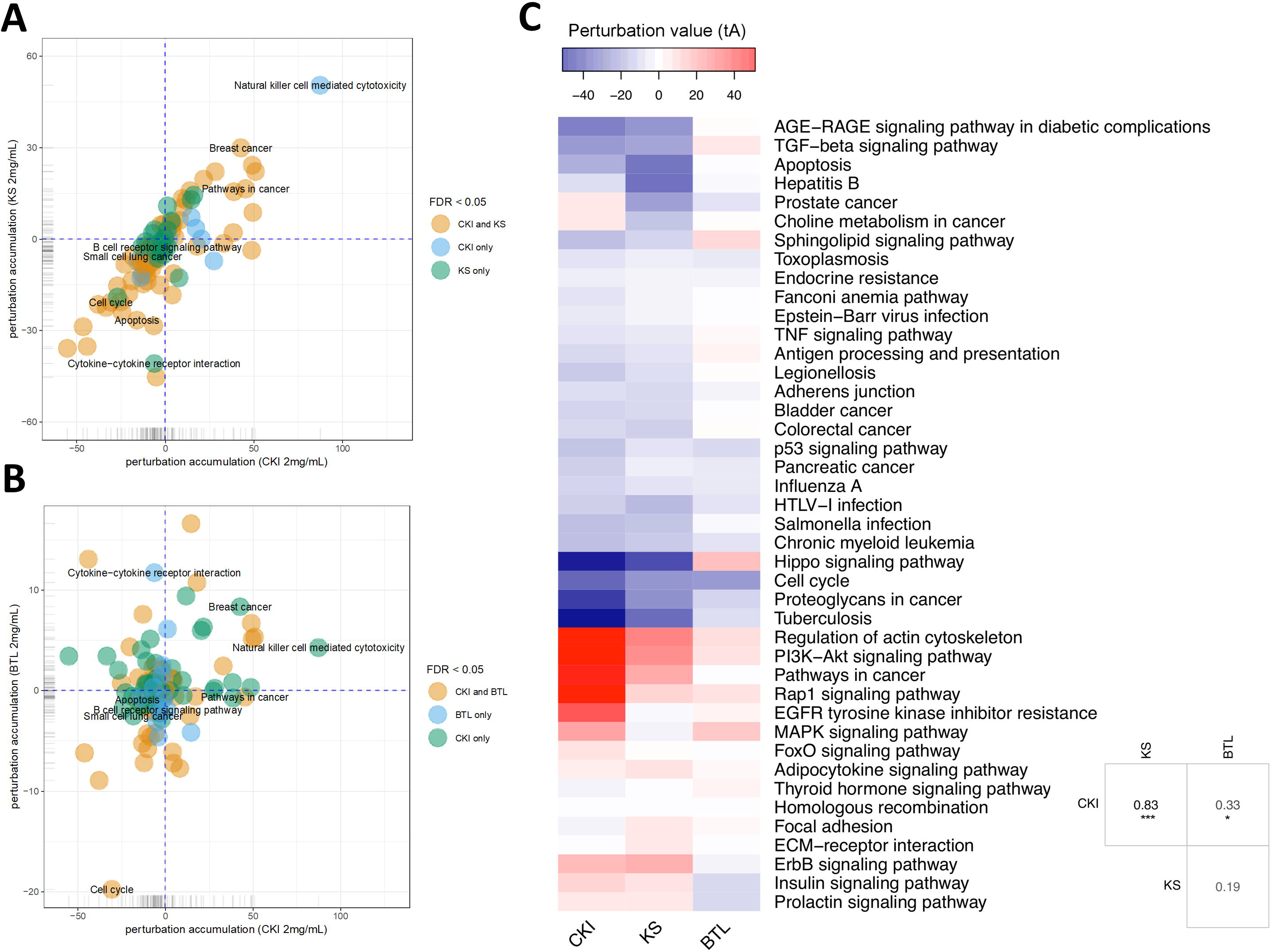
Pathway perturbation analysis for CKI and single injections on MDA-MB-231 cells. A. and B. Perturbation accumulation and significance of perturbation for different KEGG pathways treated with different injections. Positive perturbation accumulation values mean the pathway is activated and *vice versa*. Dot colors indicate whether the pathway was significantly perturbed by CKI and/or Kushen. C. Heatmap showing the perturbation value of shared significantly perturbed pathways for the three injections. Table on the right-bottom corner shows the correlation coefficients for the perturbation value for the three injections. “*” and “***” represent p<0.05 and p< 0.001 respectively (Pearson’s correlation test).

This similarity of effects between Kushen and CKI could also be seen in the high level of correlation for pathway perturbation between Kushen and CKI (0.83 correlation coefficient) compared to (0.33 correlation coefficient) for Baituling and CKI. CKI also had stronger perturbation effects on most pathways (Fig.4C). In pathways contributing to cytotoxic effects in cancer cells, such as cell cycle, p53 signaling pathway, proteoglycans in cancer and pathways in cancer, Baituling perturbed the pathways in the same direction as Kushen and seemed to reinforce those effects in CKI. However, the cytokine-cytokine receptor interaction pathway was very interesting as it was perturbed in an opposite fashion in Baituling compared to Kushen and did not show up as significantly perturbed by CKI treatment (Supplementary Fig. 2). By comparing DE genes for Baituling and Kushen in the cytokine-cytokine receptor interaction pathway (Fig.5), we observed that many genes were oppositely regulated by the two single injections, such as genes in the CXC subfamily and IL6/12-like cytokine and receptor genes, which supported the SPIA results.

**Figure 5:**
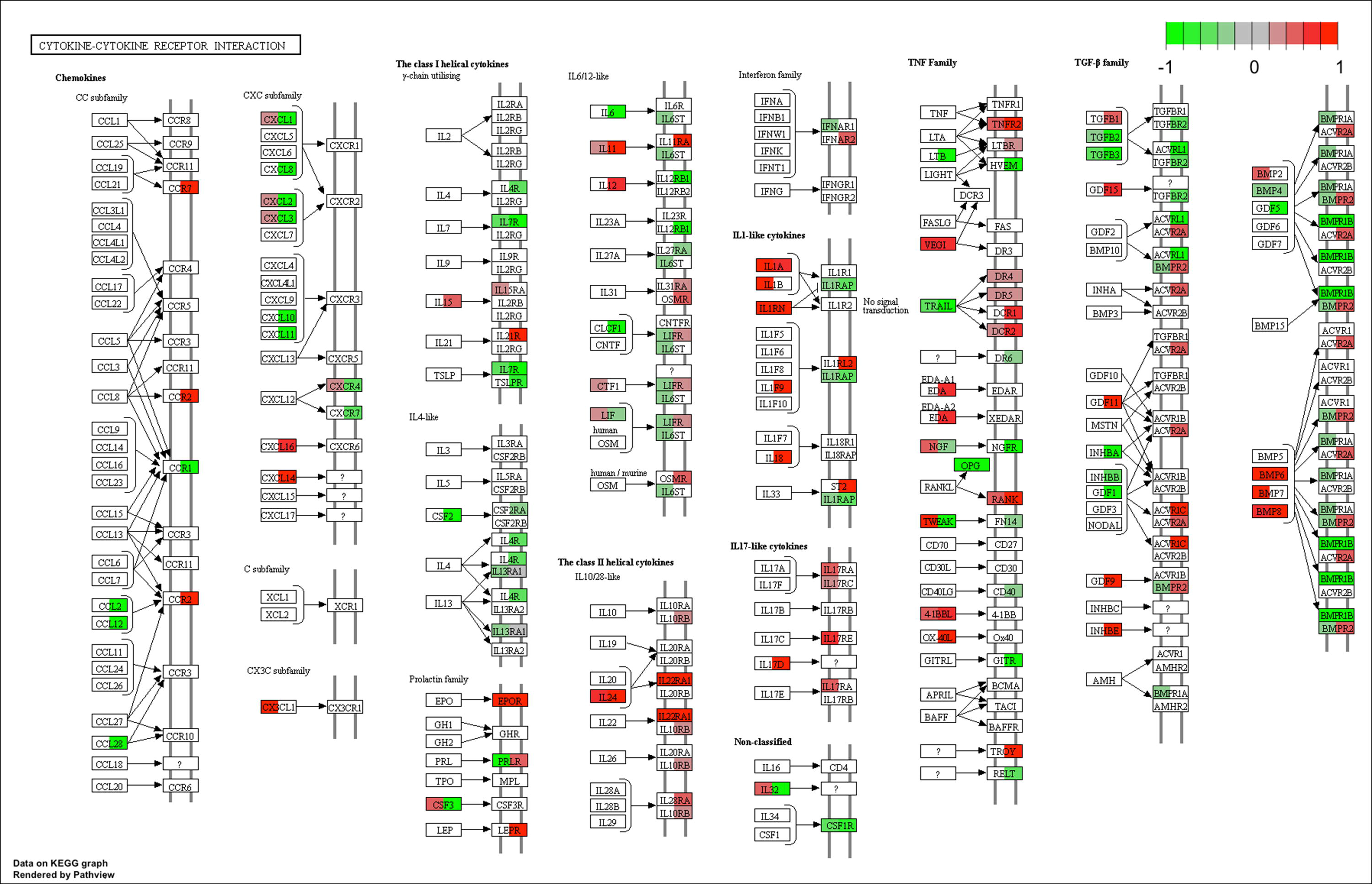
Comparison of gene expression changes caused by Baituling and Kushen treatments in the cytokine-cytokine receptor interaction pathway. Left half of each box represents the gene expression change with Baituling treatment and the right half represents the effect of Kushen treatment. White or grey colors indicate no significant change in gene expression as a function of treatment.

### The functions of Baituling in CKI

In order to investigate the function of Baituling in CKI, we identified the DE genes of CKI treatment compare to Kushen treatment. Only 308 DE genes were found (Fig.6A). KEGG analysis of these genes showed that with the exception of steroid hormone biosynthesis and transcriptional misregulation in cancer, all other pathways were related to immune function, and most genes in this set were up-regulated (Fig.6B, Supplementary Table 5). This result was consistent with findings in the SPIA analysis that Baituling tended to activate immune-related pathways. The over-represented GO terms for this gene set also included interferon-gamma production, organ or tissue specific immune response and interleukin-2 production, all aspects of immune function and which only contained up-regulated DE genes (Fig.6C).

**Figure 6:**
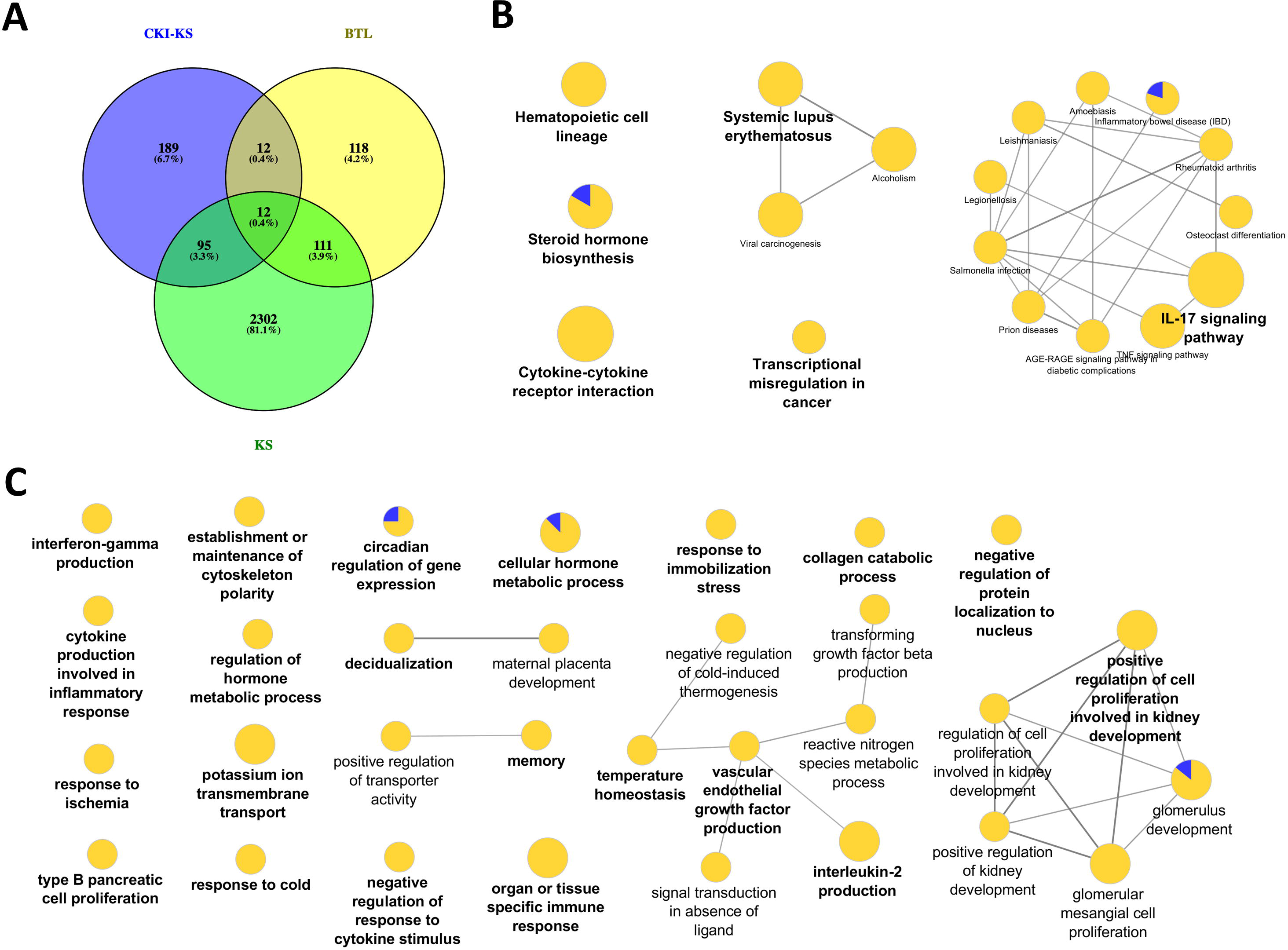
DE genes regulated by Baituling in MDA-MB-231 cells line and their functional enrichment analysis. A. Venn diagram showing the number of differentially regulated genes with CKI compared to Kushen (CKI-KS - Blue) and single herb injections (KS - Green or BTL - Yellow) compared to untreated. C. and D. Over-represented KEGG pathways and GO terms (Biological Process at 3rd level) for DE genes calculated by CKI compared to Kushen treated. Node size is proportional to the statistical significance of over-representation and colors represent the proportion of up or down-regulated genes (yellow=up-regulated and blue=down-regulated). Terms are clustered based on similar GO group (shown in bold) and related ones are connected with edges.

The genes in the common set between Kushen and CKI compared to Kushen (Fig. 6A) were originally changed with Kushen treatment and then further significantly regulated in combination with Baituling (107 genes). Only three pathways (IL-17 signaling, salmonella infection and steroid hormone biosynthesis) were over-expressed by genes in this set, indicating Baituling could also modify functions related to immune system and hormone function upregulated by Kushen in CKI (Supplementary Fig 3.). Furthermore, 24 genes appeared in both Baituling and CKI sets compared to the Kushen set (Fig. 6A), which can be regarded as the direct contribution from Baituling to the effects of CKI. Analysis of known protein-protein interactions in this gene set identified the IL-1 family and interacting proteins known to modulate immune function (Fig.7A). To verify that IL-1 protein levels were also changing as predicted, we carried out an ELISA assay to measure the IL-1 beta levels in different treatments. We were able to demonstrate that the increased IL-1 beta level in CKI treatment compared to untreated was mainly related to the effects of Baituling (Fig.7B). Altogether, these results showed that Baituling contributed to the effects of CKI primarily by altering functions related to the immune system.

**Figure 7:**
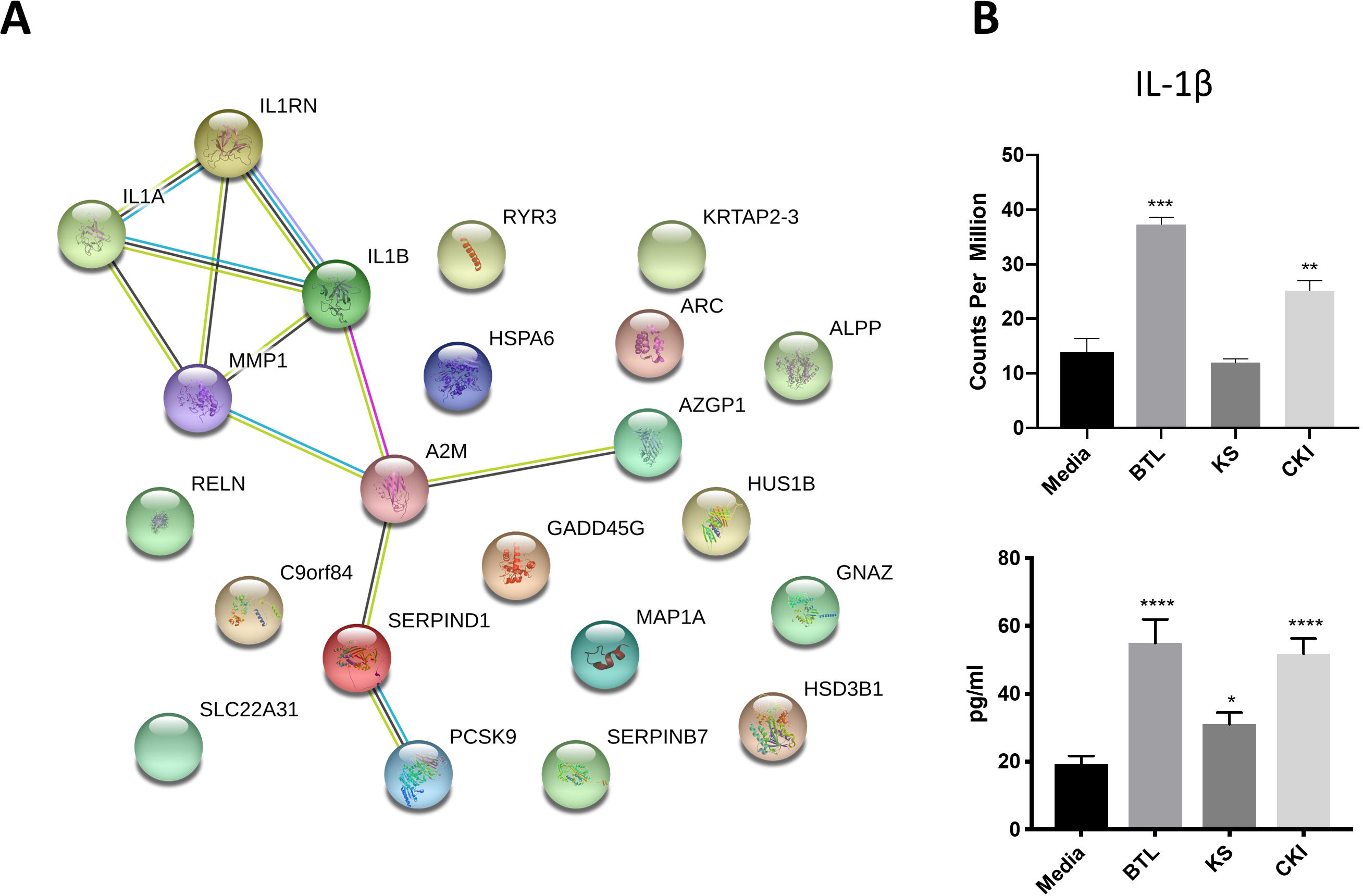
Validation of IL-1β expression changes regulated by Baituling. A. Diagram showing protein-protein interactions for common DE genes between CKI compared to Kushen, and Baituling compared to untreated. B. Comparison of expression levels and protein concentration for IL-1βwith different treatments. Top panel; IL-1β gene expression levels determined by RNA sequencing, bottom panel; levels of IL-1β in culture supernatant measured by ELISA.

In summary, we characterized the herbal compatibility of Kushen and Baitulin in CKI by comparing the individual effects of the herbal extracts to the combined extract using transcriptome analysis. In this fashion, we were able to explain the origin of CKI’s different effects in MDA-MB-231 cells. In addition, we also showed that Baituling could enhance the reduction of cell viability and increased apoptosis effects from Kushen in CKI. These results not only explained the specific molecular basis of the TCM rationale of combining Kushen with Baituling but also illustrate a general method to apply transcriptome analysis to study herbal compatibility in TCM.

## Discussion

It is undeniable that chemical composition is the basis of therapeutic effects from herbal TCM. However, because the identification and quantification of all compounds for even a single herb are still extremely difficult if not impossible, we need alternative means to conduct TCM research, particularly with respect to the study of herbal compatibility^18,19^. Herbal compatibility has a basis in TCM theory, but TCM theory is not generally accepted in Western medicine and it is difficult to map concepts from TCM theory to Western medicine. Methods that can identify the molecular consequences of TCM formulations and individual herbs can begin to provide such a map. One view of TCM is that it perturbs multiple targets or pathways with multiple low activity components to generate relatively strong effects. This is in contrast to the standard approach for pharmaceutical drug development which seeks to identify single compounds that inhibit a single pathway or target. However, this method of using one or several active compounds as representative of single herbs is problematic^10^ for understanding the roles of individual plant extracts in TCM formulations. This is illustrated by our results; although containing similar amounts of the main chemical compounds, CKI has much stronger effects than Kushen extract alone. Therefore, a formula disassembly approach that uses single herbs is a practical way to study herbal compatibility. Combined with omics techniques and network analysis, we can represent the mechanisms of TCM herbal compatibility as interactions between target networks familiar to Western medicine. In this report, we took CKI, a prescription containing only two herbs, as a proof of principle. However, related methods can easily be applied to more complex formulae.

As a frequently used herb, and because it is considered to contain the main bioactive components in CKI, Kushen or its main alkaloids and flavonoids are commonly used to represent CKI in studies^20,21^. Furthermore, TCM theory also regards Kushen as the primary herb in CKI. Our results support these hypotheses and TCM theory at different levels. First, at the chemical level, HPLC profiles showed that the source of most major components in CKI is Kushen. Second, in terms of overall efficacy, Kushen has much stronger cytotoxic effects than Baituling on various cell lines. Third, at the gene level, DE genes shared by Kushen and CKI account for 81% and 63% of total DE genes in Kushen and CKI treatments, and most of them are consistently up- or down-regulated. Furthermore, important genes are also regulated both by Kushen and CKI. Cytochrome P450 family 1 subfamily A member 1 (CYP1A1) gene, a steroid metabolizing enzyme which is important for steroid hormone responsive cancers and shown as the most over-expressed gene with CKI in our previous results, is also highly overexpressed by Kushen but not Baituling^22,23^. Also, the down-regulation of the *TP53* gene primarily results from Kushen. Our previously results based on CKI observed cells underwent apoptosis while expression of *TP53* decreased, which indicates the apoptosis induced by Kushen and CKI is not *TP53*-dependent^11^. Finally, GO and KEGG over-representation analysis also indicated that the genes in major cancer related pathways and terms perturbed by CKI including cell cycle, DNA replication, and cell migration, are induced by the shared DE gene set between Kushen and CKI. Taken together, our findings agree with TCM theory which considers Kushen as the principle herb in CKI, and they map this specific part of TCM theory to effects on specific genetic networks and pathways.

On the other hand, we could find no Baituling literature and studies on macrozamin to support its application in CKI^24^. From our HPLC and cytotoxic assay results, Baituling injection does not contain many components and has no significant effects on cancer cells. In addition, NAseq analysis of Baituling-treated samples are close to the untreated samples in the multidimensional scaling plot, and only 253 DE genes compared to untreated were detected. However, after comparing the differences between CKI and Kushen, we found Baituling has a strong reinforcing effects on Kushen. The SPIA results showed Baituling can enhance many pathways’ perturbation strength compared to Kushen treatment, including cell cycle, pathways in cancer and proteoglycans in cancer. In addition, many DE genes in immune-related pathways and GO terms are over-represented in CKI compared to Kushen. Together with the opposing direction of perturbation for the cytokine-cytokine receptor interaction pathway caused by Kushen and Baituling, we can conclude Baituling may also contribute to the immune regulatory effects of CKI. This was confirmed by our measurements of IL-1**β**, which was significantly up-regulated by Baituling and CKI. In conclusion, our results indicate that Baituling, an adjuvant herb in CKI according to TCM theory, may enhance the anticancer effects of Kushen and contribute to immune regulation.

In conclusion, we explained the TCM herbal compatibility of CKI in the context of pathway perturbations using transcriptome analysis. Kushen primarily contributes to CKI effects on cancer cells by perturbing cell cycle regulation and other functions^11,14^, meanwhile, Baituling enhanced potential anticancer effects for Kushen and activated the immune system. Therefore, the two herbs in CKI collaborate with differences in effects, which is very similar to formulation theory in TCM. Compared to previous studies on herbal compatibility, our method can explain the beneficial interaction pattern of herbs in TCM formulae in a more systematic and comprehensive fashion.

## Methods

### Cell culture and drugs

MDA-MB-231, HepG2 and A431 cells were purchased from ATCC (VA, USA). All cell lines were cultured at 37°C with 5% CO_2_ in DMEM (Thermo Fisher Scientific, MA, USA) with 10% fetal bovine serum (Thermo Fisher Scientific). CKI, Baituling, and Kushen injections were provided by Zhendong pharmaceutical Co.Ltd (China). Baituling and Kushen injections were manufactured using the same processes as CKI and diluted to the equivalent concentration of CKI (total alkaloid concentration at 2 mg/ml).

All *in vitro* assays were conducted in 6-well or 96-well plates. The seeding density was 4 × 10^5^ for 6-well plates across all three cell lines. For 96-well plates, MDA-MB-231 cells, A431 cells and HepG2 cells were seeded at 1.6 × 10^5^ cells/well, 8 × 10^4^ cells/well and 4 × 10^3^ cells/well respectively. Cells were cultured overnight before drug treatment and the treatment time was 48 hours for all assays.

### Components comparison with HPLC

CKI, Kushen or Baituling injection was diluted 1:20 with MilliQ water and then analyzed on a photodiode-array UV-Vis detector equipped Shimadzu HPLC (Japan) with a preparative C18 column (5 μm, 250 × 10 mm, Phenomenex, CA, USA). The recording range is from 200 nm to 280 nm, with monitoring at 215 nm. 0.01M ammonium acetate (adjusted to pH 8.0, solvent A) and acetonitrile + 0.09 % trifluoroacetic acid (solvent B) were used as mobile phase and flow rate is 2 ml/min with linear gradient elution (0 min, 100 % A; 60 min, 65 % A, 70 min, 100 % A). Nine Standard compounds, including Oxymatrine, Oxysophocarpine, N-methylcytisine, Matrine, Sophocarpine, Trifolirhizin, Adenine, Sophoridine (Beina Biotechnology Institute Co., Ltd, China), and macrozamin (Zhendong Pharmaceutical Co.Ltd, China), were used to characterize peaks in the HPLC profile.

### Cell viability assay

Cells were cultured and treated in 96-well plates. After 48 hours drug treatment, 50 μl of XTT:PMS (at 1 mg/ml and 1.25 mM, respectively, and combined at 50:1 ratio, Sigma-Aldrich, MO, USA) was added into each well and incubated 4 hours for the measurement of cell viability. A Biotrack II microplate reader was used to detect the absorbance at 492 nm.

### Apoptosis rate with cell cycle assay

After treatment, cells were harvested from 6-well plates and stained with propidium iodide (PI; Sigma-Aldrich) as previously described^25^. The stained cells were quantified on a BD LSR Fortessa-X20 (BD Biosciences, NJ, USA) and the data were analyzed with FlowJo software (TreeStar Inc., OR, USA).

### qPCR for transcriptome validation

The assay was performed as previously described^11^. The sequences of all primers are shown in the Supplementary Table.

### RNA extraction and sequencing

After treatment with injections, MDA-MB-231 cells were harvested from 6-well plates and snap-frozen with liquid nitrogen. Total RNA was isolated using the PureLink RNA mini kit (Thermo Fisher Scientific). Quality and quantity of RNAs were measured with a Bioanalyzer at the Cancer Genome Facility of the Australian Cancer Research Foundation (Australia) to ensure RIN>7.0 and sent to Novogene (China) for sequencing with paired-end 150 bp reads on an Illumina HiSeq X platform.

### Transcriptome data analysis

The adaptors and low-quality sequences in raw reads were trimmed with Trim_galore (v0.3.7, Babraham Bioinformatics) using parameters: --stringency 5 --paired. STAR (v2.5.3a) was used to align reads to reference genome (hg19, UCSC) with parameters: --outSAMstrandField intronMotif --outSAMattributes All --outFilterMismatchNmax 10 --seedSearchStartLmax 30^26^. Differentially expressed genes were calculated with edgeR (v3.22.3) and selected with false discovery rate (FDR) < 0.05 and Log fold change >1 or <−1^17^.

The GO and KEGG over-representation analyses were performed with ClueGO and visualized with Cytoscape v3.6.0 with following parameters: right-sided hypergeometric test for enrichment analysis; p values were corrected for multiple testing according to the Benjamini-Hochberg method and biological process at 3rd level for GO terms^27,28^. The Signalling Pathway Impact Analysis (SPIA) package in R was used to conduct the pathway perturbation analysis using all DE genes(FDR<0.05)^29^. R Pathview package was used to visualize specific KEGG pathways^30^. String (V11.0) was used to identify protein-protein interactions with a threshold of 0.4 for minimum interaction score^31^.

### ELISA for IL-1β level

A431 cells were treated with different injections for 48 hours in 96-well plates and the cell culture supernatant was collected and tested for the level of IL-1β by ELISA using human interleukin-1 beta ELISA kit (Biosensis, CA, USA) according to the kit protocol. The absorbance at 450nm was detected with Multiskan Ascent Plate Reader.

## Supporting information

Legends for Supplementary Material

