## Supplementary material for "Understanding the Mechanistic Contribution of Herbal Extracts in Compound Kushen Injection with Transcriptome Analysis": Legends for Supplementary Material

Supplementary Figure 1: Multiple dimensional scaling (MDS) plot for samples based on expression profiles of all genes (Untreated in black, Baituling in red, Kushen in green and CKI in blue).

Supplementary Figure 2: Heatmap showing the perturbation value of significantly perturbed pathways only for one or two injections. The significantly perturbed pathways were selected with criteria FDR < 0.05.

Supplementary Figure 3: Over-represented KEGG pathways and their contained genes showing shared DE genes between CKI (DE calculated by comparison to Kushen treated) and Kushen (DE calculated by comparison to untreated). Node size is proportional to the statistical significance of over-representation and genes are connected to their belonged pathways with edges.

Supplementary Table 1: Mapping rates for each RNA-seq result.

Supplementary Table 2: Primer sequences for RT-qPCR.

Supplementary Table 3: DE gene lists for different comparisons.

Supplementary Table 4: SPIA results of significantly perturbed KEGG pathways for three injections.

Supplementary Table 5: Significantly over-represented GO and KEGG terms (count > 4 and P-value < 0.05)

i. Sheet 1-2: GO and KEGG enrichment of DE genes shared by CKI compared to ‘untreated’ and Kushen compared to ‘untreated’.
ii. Sheet 3-4: GO and KEGG enrichment of DE genes for CKI compared to Kushen.
