## Supplementary figures and images for "Understanding the Mechanistic Contribution of Herbal Extracts in Compound Kushen Injection with Transcriptome Analysis"

### Supplementaray Figure 1

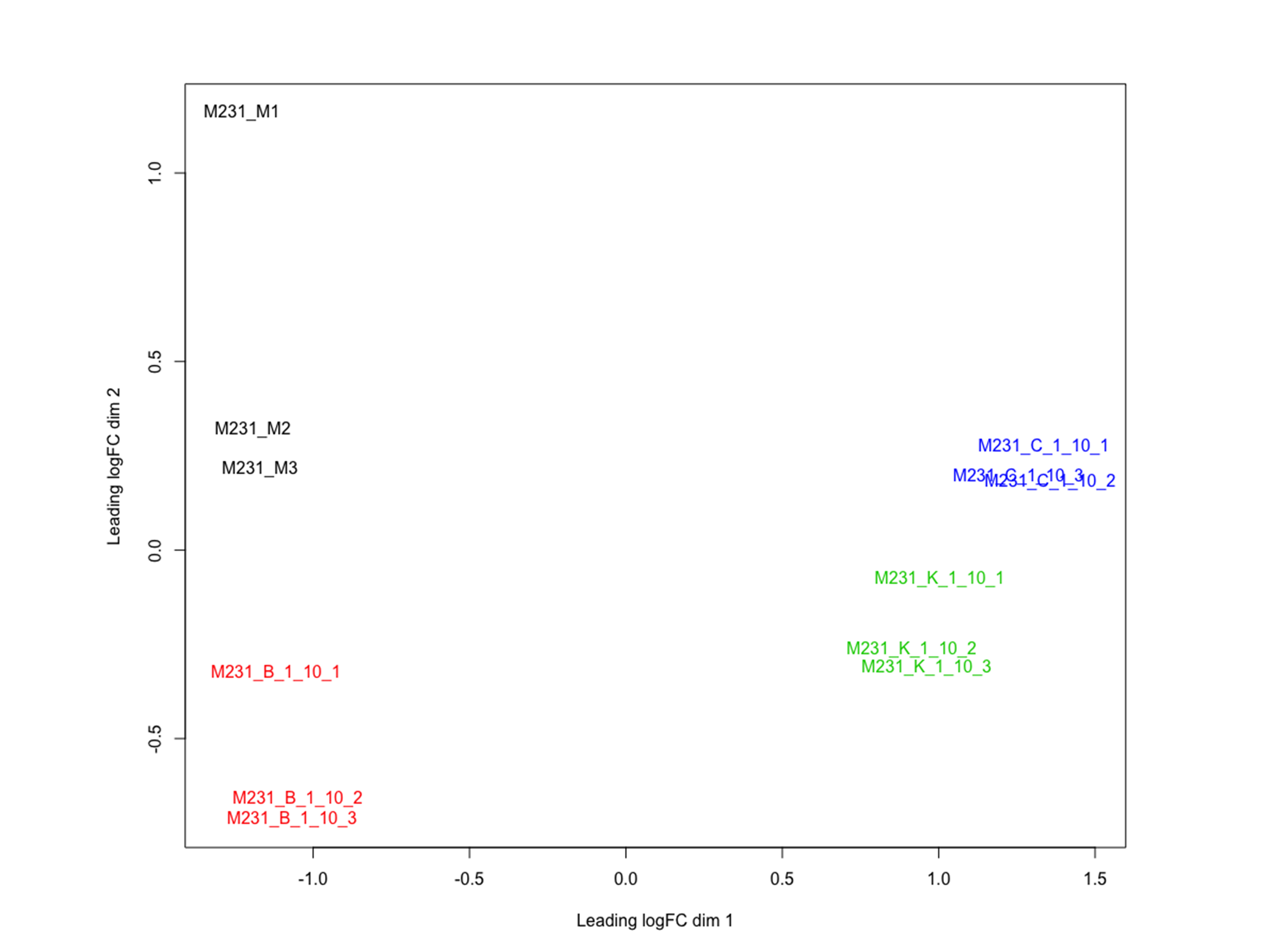

### Supplementary Figure 2

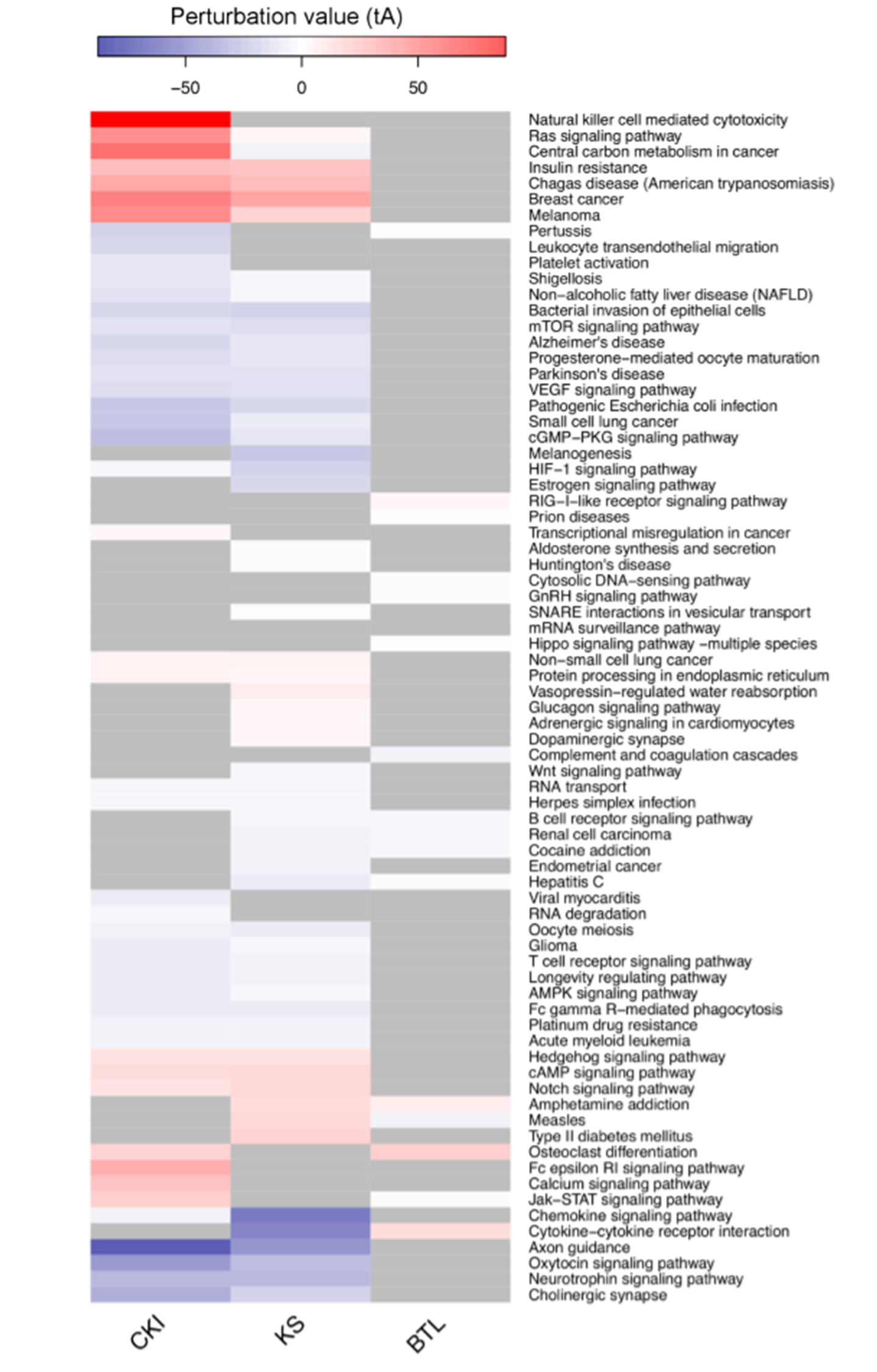

### Supplementary Figure 3

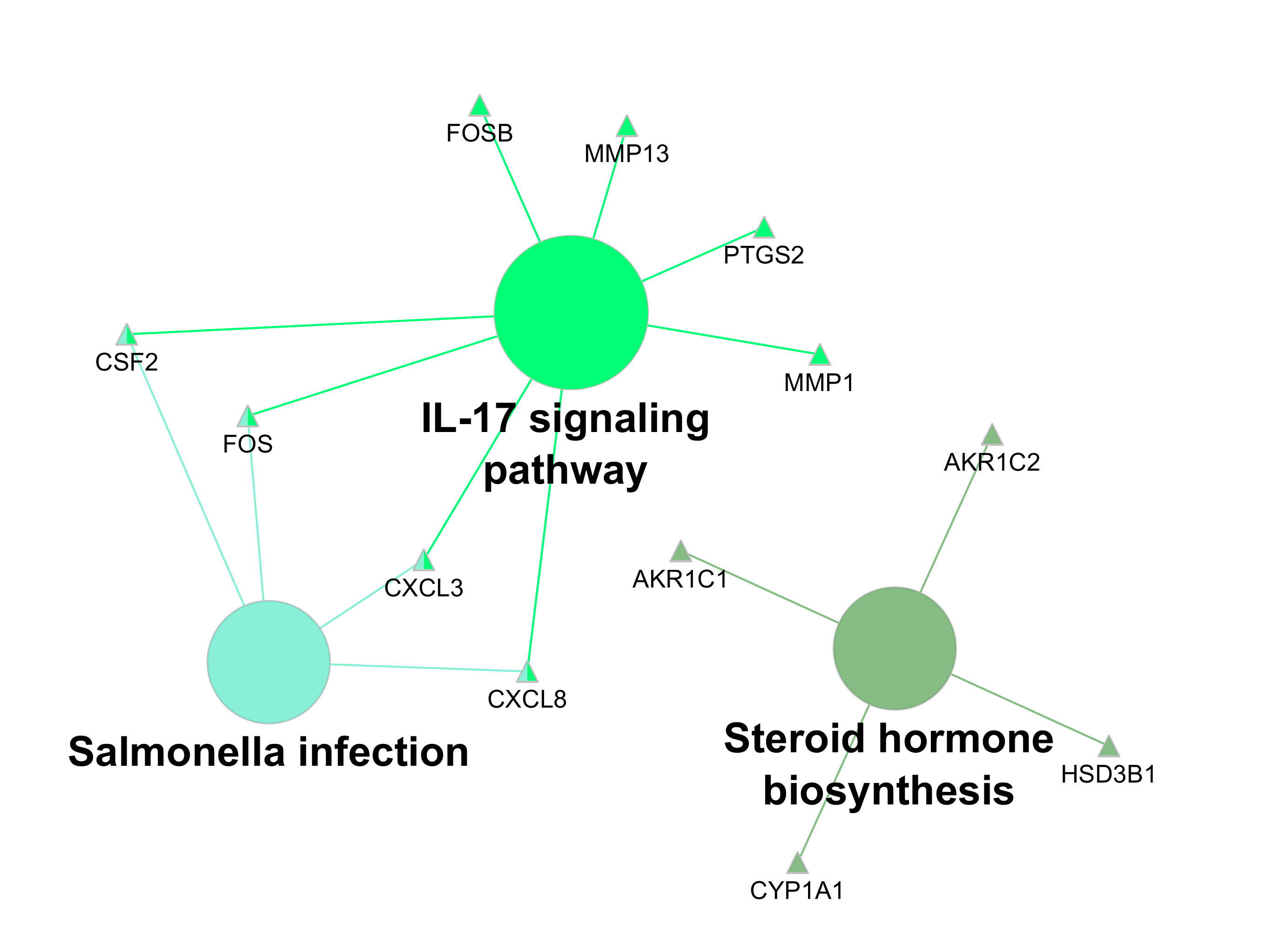
